# Local Orchestration of Global Functional Patterns Supporting Loss and Restoration of Consciousness in the Primate Brain

**DOI:** 10.1101/2023.06.30.547281

**Authors:** Andrea I. Luppi, Lynn Uhrig, Jordy Tasserie, Camilo M. Signorelli, Emmanuel Stamatakis, Alain Destexhe, Bechir Jarraya, Rodrigo Cofre

## Abstract

A central challenge of neuroscience is to elucidate how the orchestration of brain function is modulated by different states of consciousness. Here, we investigate the link between distributed structural and functional brain organisation in functional MRI signals of non-human primates, through bi-directional causal manipulations of consciousness. During varying levels of propofol, sevoflurane, or ketamine anaesthesia, and subsequent restoration of responsiveness by deep brain stimulation of the central thalamus, we investigate how loss of consciousness impacts distributed patterns of structure-function organisation across scales. Combining the specificity of electrical stimulation with global fMRI coverage of the entire cortex, we report that distributed brain activity under anaesthesia is increasingly constrained by brain structure across scales, coinciding with anaesthetic-induced collapse of multiple dimensions of hierarchical cortical organisation. Crucially, we show that these distributed signatures of anaesthetic-induced loss of consciousness are observed across different anaesthetics, and they are reversed by electrical stimulation of the central thalamus, coinciding with recovery of behavioural markers of consciousness during propofol anaesthesia. No such effects were observed upon stimulation of a control anatomical site, ventral lateral thalamus, demonstrating specificity. Through causal manipulations of consciousness that integrate pharmacology and electrical intracranial stimulation of the thalamus, our results identify global signatures of consciousness that are under local causal control by specific nuclei of the thalamus. Overall, the present work broadens our understanding of the link between brain network organisation and distributed function in supporting consciousness, and the interplay between local and global functional architecture.

## Introduction

Despite significant progress in recent years, understanding how the activity and connectivity of the brain support and causally determine different states of consciousness remains a major challenge for modern neuroscience (Dehaene, Lau and Kouider, 2017; Kringelbach and Deco, 2020). The traditional approach to the neuroscience of consciousness has been to look for specific and localised brain regions that support consciousness (Crick and Koch, 1998; Seth and Bayne, 2022). However, there is now increasingly compelling evidence for a role of distributed, large-scale patterns of cortical organisation supporting cognitive function, dysfunction, (Margulies *et al*., 2016; Huntenburg, Bazin and Margulies, 2018; Fulcher *et al*., 2019; Paquola *et al*., 2019; Bethlehem *et al*., 2020; Cross *et al*., 2021) and consciousness (Dehaene and Naccache, 2001; Dehaene and Changeux, 2011; Atasoy *et al*., 2017; Atasoy, Deco, *et al*., 2018). Recent work has demonstrated that multiple pathological and pharmacological perturbations of consciousness induce consistent reorganisation of the brain’s functional architecture along functional and anatomical axes of cortical organisation (Huang, Mashour and Hudetz, 2023; Li *et al*., 2023; Luppi, Hansen, *et al*., 2023).

This distributed approach goes beyond viewing the brain in terms of brain regions and fixed networks, emphasising the role of dynamics and functional organisation. For instance, when consciousness is lost, the functional interactions between brain regions (“functional connectivity”) become increasingly similar to the pattern of direct anatomical connections between regions (“structural connectivity”) (Barttfeld *et al*., 2015; Ma, Hamilton and Zhang, 2017; Uhrig *et al*., 2018; Demertzi *et al*., 2019; Gutierrez-Barragan *et al*., 2022; Tasserie *et al*., 2022). More recently, evidence has shown that the organisation of distributed brain function is perturbed in a scale-specific manner under several pharmacological and pathological perturbations of consciousness (Huang, Mashour and Hudetz, 2023; Luppi, Vohryzek, *et al*., 2023) – with anaesthesia and disorders of consciousness increasing the dependence of brain function on structural connectivity across scales.

Here, we seek to integrate the localised and distributed approaches to characterise the functional architecture of the macaque brain. To this end, we leverage functional MRI (fMRI) data from non-human primates in the awake state, under loss of consciousness induced by three different anaesthetics (sevoflurane, propofol, ketamine) and restoration of consciousness by deep brain stimulation (DBS). Specifically, we consider the brain’s hierarchical organisation across scales, by studying its principal gradient of functional connectivity, and its complex relationship with the other functional eigenmodes of the brain. We then integrate brain structure and function, by decomposing brain activity into distributed patterns of structure-function coupling, termed structural eigenmodes. Through this decomposition, we quantify the extent to which brain activity is constrained by the underlying network of structural connectivity across multiple spatial scales, from large-scale to localised.

Comparing the brain-wide effects of different anaesthetics enables us to disentangle which aspects of the brain’s functional organisation support consciousness, being consistently targeted by anaesthetics, despite their distinct molecular mechanisms. Crucially, we also aim to obtain more stringent evidence for the causal relevance of distributed signature of consciousness, by determining whether the reorganisation of distributed brain function consistently induced by different anaesthetics can be reversed by targeted stimulation of different subregions of the thalamus, a brain structure that has been repeatedly associated with supporting consciousness (Tononi, 2004; Schiff *et al*., 2007; Staunton, 2008; Mhuircheartaigh *et al*., 2010; Lioudyno *et al*., 2013; Ní Mhuircheartaigh *et al*., 2013; Vijayan *et al*., 2013; Akeju *et al*., 2014; Lewis *et al*., 2015; Flores *et al*., 2017; Hemmings *et al*., 2019; Kelz *et al*., 2019; Zucca *et al*., 2019; Mashour *et al*., 2020; Redinbaugh *et al*., 2020, 2022; Afrasiabi *et al*., 2021; Bastos *et al*., 2021; Setzer *et al*., 2022; Gammel *et al*., 2023; Kantonen *et al*., 2023). We do so, using a complementary but independently acquired dataset of fMRI in macaques recorded in the awake state and during anaesthesia with and without concurrent deep brain stimulation of the centro-medial thalamus (CT) or ventro-lateral thalamus (VT), previously shown to induce recovery of responsiveness according to the location and intensity of the electrical input (Tasserie *et al*., 2022). This dual causal manipulation – pharmacology and electrical stimulation – provides us with a unique opportunity to study functional changes that are observed during loss of consciousness and reappear upon recovery of consciousness, despite continuous anaesthetic infusion. By combining the broad spatial coverage of fMRI with stimulation of specific nuclei of the thalamus, we can consider both local and global effects. Through this approach, we show that distributed cortical signatures of consciousness are under local control specifically by the central thalamus, thereby integrating the distributed and localised perspectives.

## Results

Here, we studied distributed functional patterns in the macaque cerebral cortex. Specifically, we measure how these patterns are reorganised under different anaesthetic agents and by direct stimulation of different thalamic nuclei.

### Anaesthetic-Induced Collapse of the Principal Functional Gradient

Our first approach to study distributed brain function and its alteration under anaesthesia is through the lens of the principal gradient of functional connectivity. Functional gradients, as the eigenmodes of the brain’s functional connectivity, provide a low-dimensional representation of the similarity between different regions’ patterns of functional connectivity. Along each dimension (represented by an eigenmode), regions whose connectivity with the rest of the brain is more similar will exhibit more similar values – whose spatial variation delineates the gradient. In particular, the principal functional gradient (eigenmode associated with the principal eigenvalue) captures the direction of maximal spatial variation in functional organisation: its range can be interpreted as reflecting the distance between the two extremes of the cortical processing hierarchy, reflecting the depth of information processing. This principal gradient can also be reshaped by pharmacological intervention (Girn *et al*., 2022; Timmermann *et al*., 2023), rendering this approach a promising tool for the present study. Here, we hypothesise that this depth should be reduced in the unconscious brain, when processing of information is manifestly impaired. Indeed, such an observation was very recently reported in humans, with different gradients of functional connectivity collapsing as a result of different pharmacological and pathological perturbations, including anaesthesia (Huang, Mashour and Hudetz, 2023).

Here, we delineate gradients using the common nonlinear dimensionality reduction technique known as diffusion map embedding (see Methods). We observed anaesthetic-induced collapse of the principal gradient of macaque functional connectivity, which is restored by centro-median thalamic stimulation at high amplitude. The range of the principal gradient of macaque functional connectivity is significantly reduced by anaesthesia, regardless of the specific agent used (Figure 1 and Figure S1), and is significantly increased back to awake levels by low amplitude stimulation of the centro-median thalamus (Figure 2 and Figure S1). In general, the second gradient appears less sensitive to perturbations, both in terms of reflecting the effect of anaesthetics, and in terms of reflecting restoration due to CT stimulation (Figure S2). Surprisingly, a weaker effect was observed when stimulation at the same location at high amplitude, although both induce significant restoration of the principal functional gradient, compared with stimulation of the control site in the ventral-lateral thalamus.

**Figure 1.**
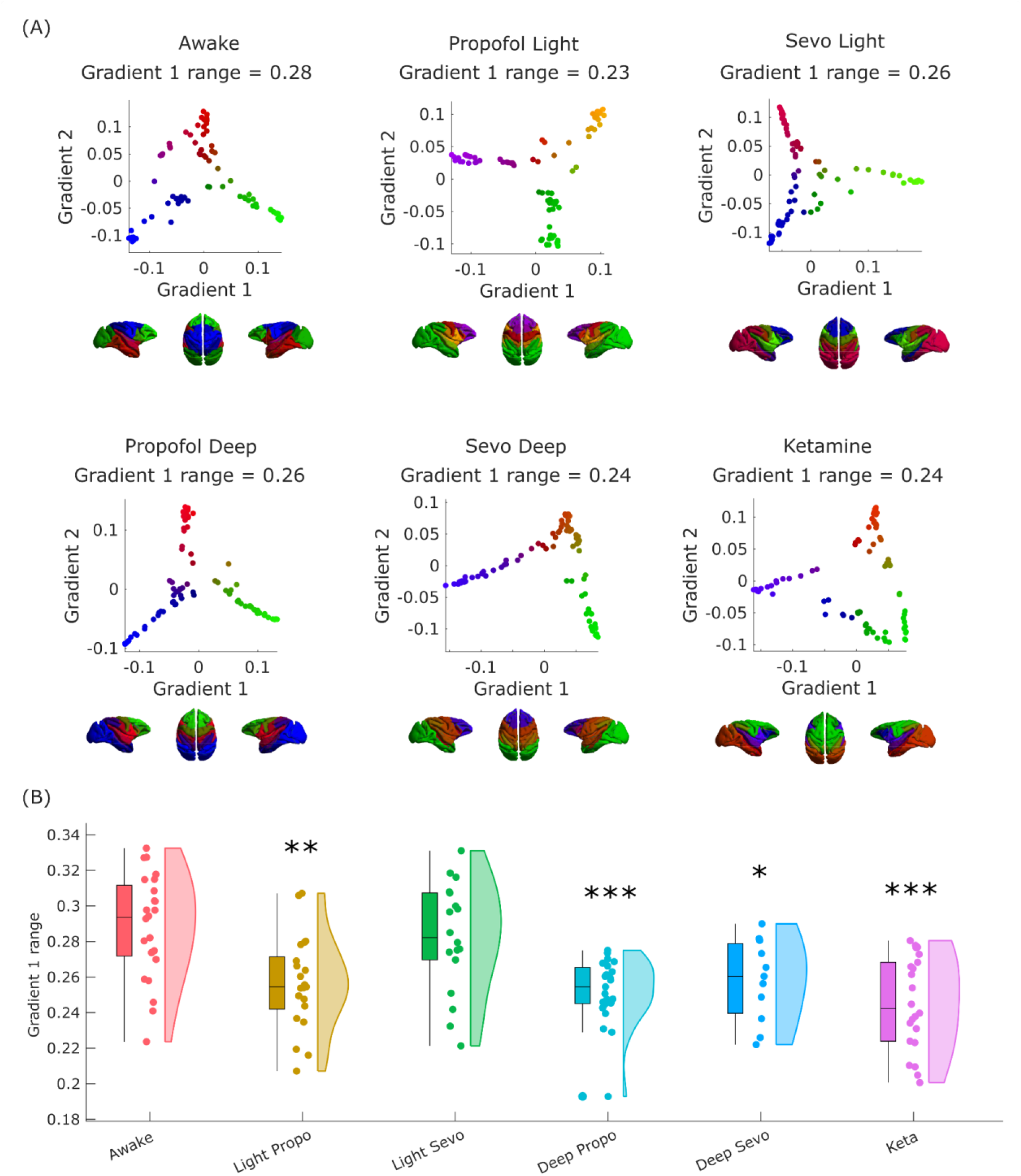
Anaesthetic-induced collapse of the principal gradient of macaque functional connectivity. (A) Scatter plots show the first two principal gradients of macaque functional connectivity (obtained from diffusion graph embedding: see Methods) for the group-averaged FC matrix of the awake condition, and each anaesthetised condition. The gradients are also plotted on the cortical surface of the macaques, with colour representing the position of each region along each gradient. (B) The range of the principal gradient of macaque functional connectivity across wakefulness and different anaesthetic conditions. Box plots indicate the median and interquartile range of the distribution. * *p* < 0.05; ** *p* < 0.01; *** *p* < 0.001 (FDR-corrected), compared against Awake condition; see Figure S1 for comparisons between all pairs; see Figure S2 for gradient 2 results).

**Figure 2.**
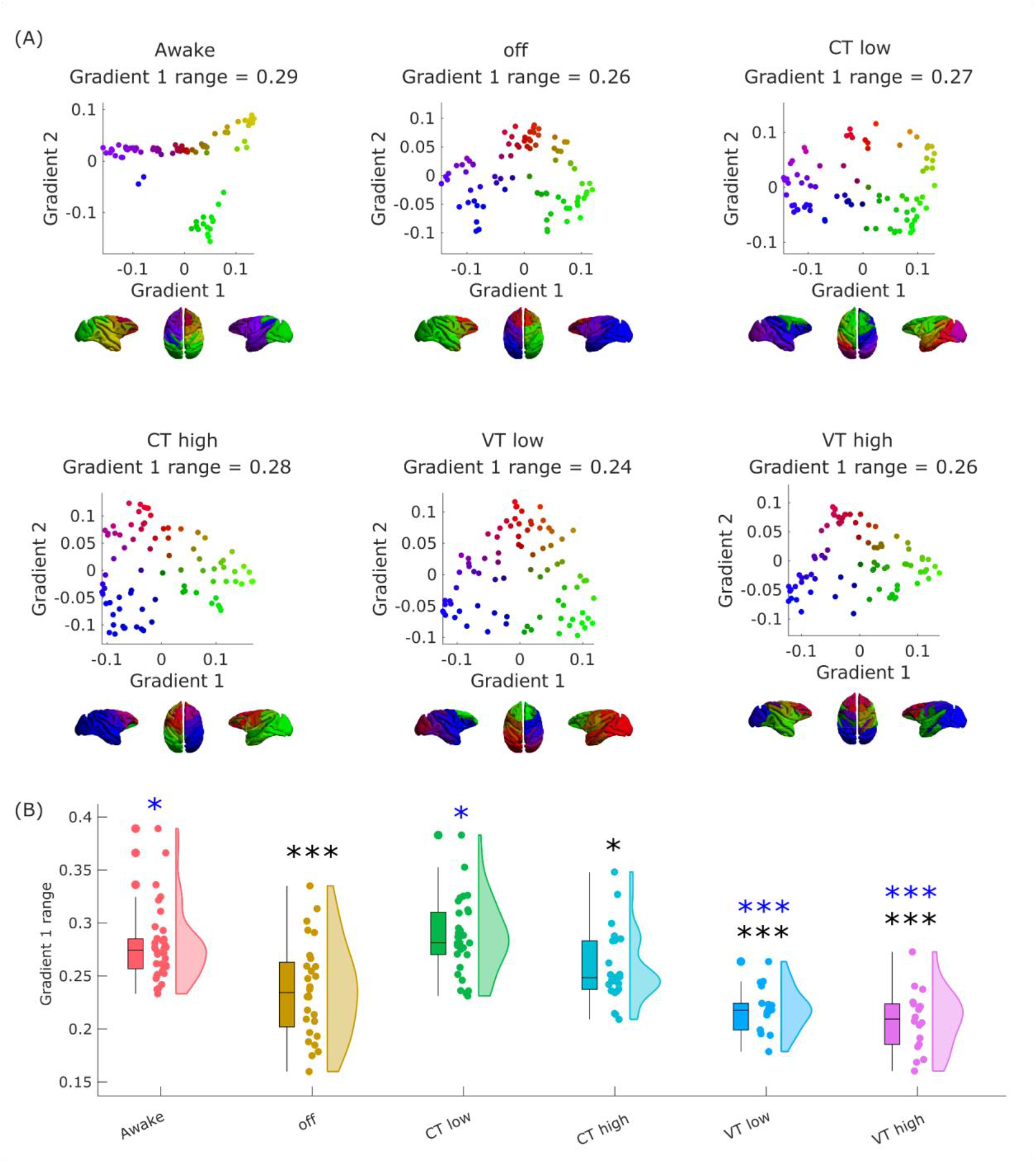
Anaesthetic-induced collapse of the principal gradient of macaque functional connectivity is restored by thalamic stimulation. (A) Scatter plots showing the first two principal gradients of macaque functional connectivity (obtained from diffusion graph embedding) for the group-averaged FC matrix of the awake condition, and of the anaesthetized state, with and without DBS thalamic stimulation. The gradients are also plotted on the cortical surface of the macaques, with colour representing the position of each region along each gradient. (B) Effect of thalamic deep-brain stimulation on the range of the principal gradient of functional connectivity in macaques during anaesthesia. Box plots indicate the median and interquartile range of the distribution. * *p* < 0.05; ** *p* < 0.01; *** *p* < 0.001 (FDR-corrected), compared against Awake condition; * *p* < 0.05; ** *p* < 0.01; *** *p* < 0.001 (FDR-corrected), compared against CT high condition; see Figure S1 for comparisons between all pairs; see Figure S2 for gradient 2 results).

### Hierarchical Integration Across Scales is Compromised under Anaesthesia

Whereas the range of the principal functional gradient reflects the putative depth of the information-processing hierarchy, hierarchical organisation can also manifest in terms of nested relationships between the system’s parts at different scales: lower elements in the hierarchy are recursively combined to form the higher elements (Hilgetag and Goulas, 2020). This perspective makes it possible to consider the interplay of global integration and local segregation of brain signals – which is a central feature of prominent scientific accounts of consciousness. Classical graph-theoretic measures such as small-worldness and modularity quantify integration and segregation at a single scale, making them inadequate to capture these properties across multiple hierarchical modules (Newman, 2006; Rubinov and Sporns, 2010; Wang *et al*., 2021). Instead, we can obtain insight into the hierarchical relationships between different scales of distributed activation in the brain by going beyond the principal gradient alone, and instead considering all functional eigenmodes (Wang *et al*., 2021). This is achieved by characterising the similarities and differences of regions’ allegiance across eigenmode scales, resulting in a nested, modular structure that identifies a hierarchical sub-division of the functional connectome into nested modules (see Figure 3). Hierarchical integration and segregation can then be quantified by the relative prevalence of the different eigenmodes, indicated by their associated eigenvalues (Wang *et al*., 2021).

**Figure 3.**
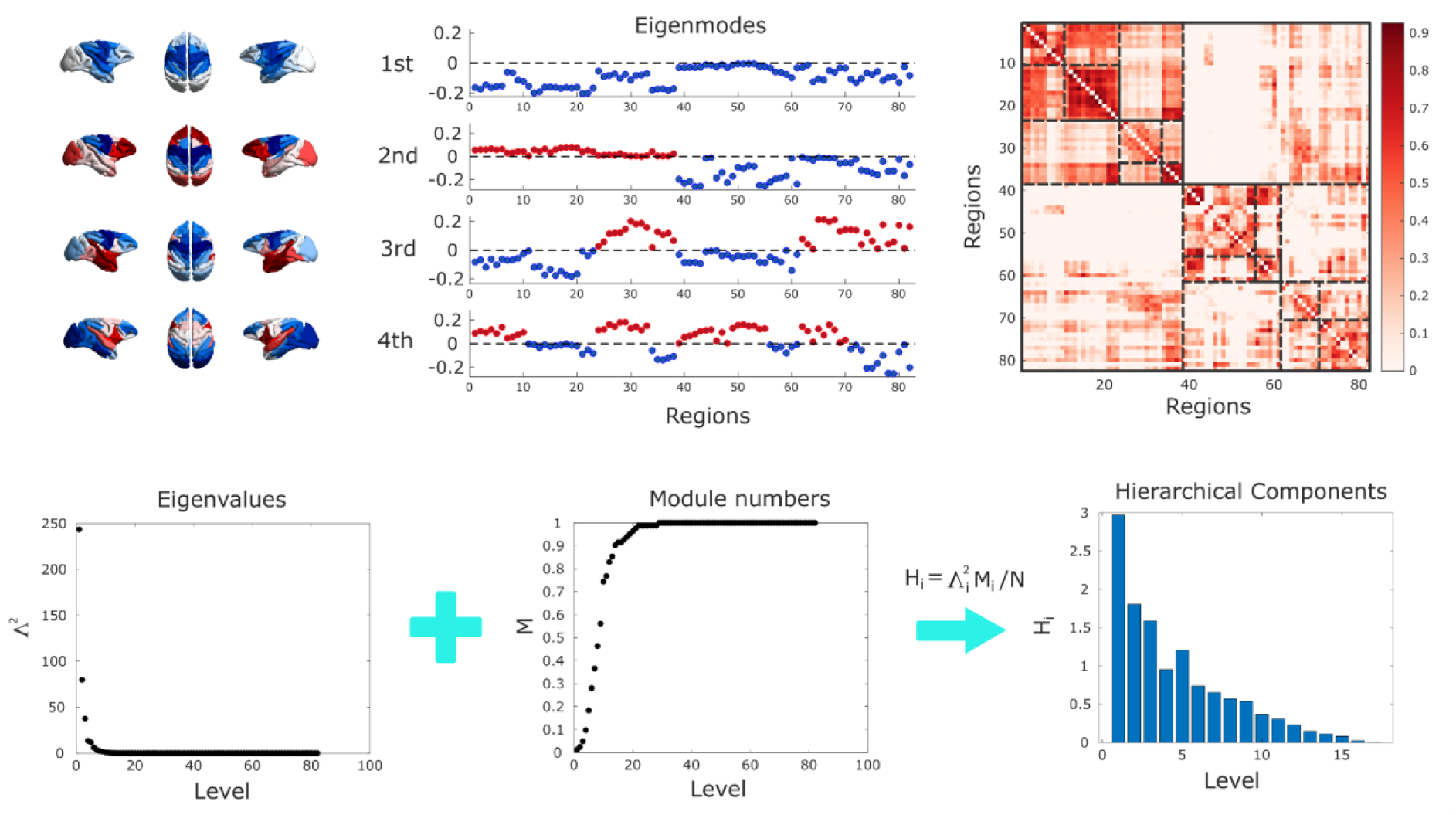
Hierarchical integration quantified from brain functional eigenmodes. Top: Each eigenmode (except the first) divides cortical regions into two groups, at progressively finer scales. Combining the different groupings identifies a hierarchical sub-division of the functional connectome into nested modules. Bottom: The relative weight of each eigenmode is given by its associated eigenvalue. The first eigenmode corresponds to global integration, and segregation is then reflected by the contribution of the other eigenmodes.

We observe that the hierarchical brain integration of macaque eigenmodes is reshaped by loss and recovery of consciousness. Eigenmode-based hierarchical integration is significantly reduced in the macaque brain under anaesthesia, whether induced by sevoflurane, propofol, or ketamine (Figure 4 and Figure S3). In contrast, hierarchical segregation is nearly insensitive to the effects of anaesthesia (Figure S4). The reduction in eigenmode-based hierarchical integration induced by anaesthesia is also specifically restored by high amplitude electrical stimulation delivered in the centro-median nucleus of the thalamus, bringing it back to the level of the awake state (Figure 4). These results reveal novel signatures of consciousness restored by deep brain stimulation of the central thalamus under anaesthesia.

**Figure 4.**
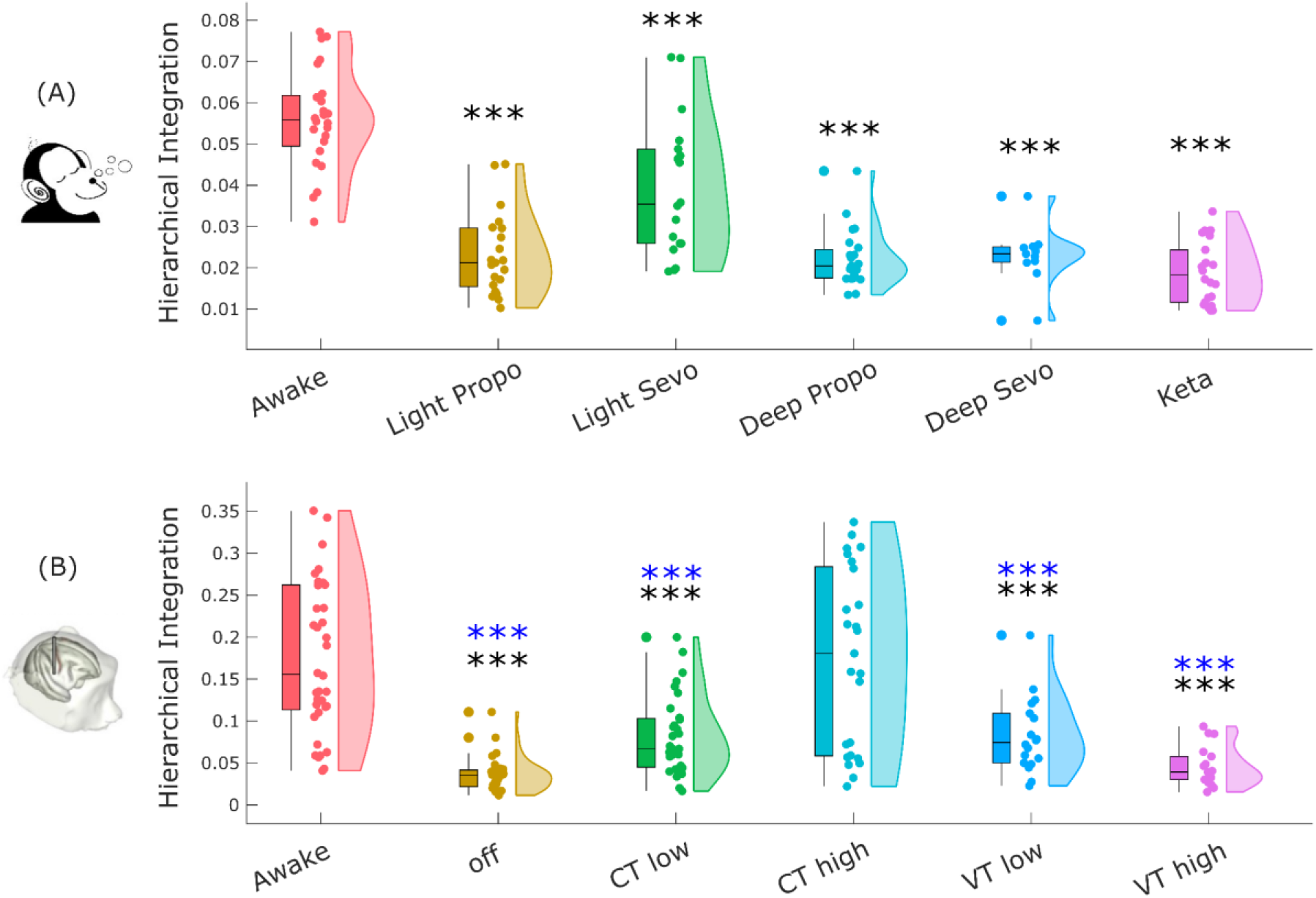
Hierarchical brain integration from macaque eigenmodes is reshaped by loss and recovery of consciousness. (A) Eigenmode-based hierarchical integration is significantly reduced in the macaque brain under anaesthesia, whether induced by sevoflurane, propofol, or ketamine. (B) The reduction in eigenmode-based hierarchical integration induced by anaesthesia is restored to awake levels by electrical stimulation of the centro-median thalamus with high (5V) current. Box plots indicate the median and interquartile range of the distribution. * *p* < 0.05; ** *p* < 0.01; *** *p* < 0.001 (FDR-corrected), compared against Awake condition; * *p* < 0.05; ** *p* < 0.01; *** *p* < 0.001 (FDR-corrected), compared against CT high condition; see Figure S3 for comparisons between all pairs, and Figure S4 for hierarchical segregation).

### Harmonic Mode Decomposition Reveals Increased Structural Constraints on Brain Activity

Functional brain activity and connectivity unfold over the network of physical white matter pathways between brain regions: the structural connectome. Therefore, for our final investigation we go beyond function alone, by explicitly integrating brain activity and structural connectivity. To this end, inspired by the work of Atasoy and colleagues (2016) in humans, we leverage the mathematical framework of “harmonic mode decomposition” (Atasoy, Donnelly and Pearson, 2016), which decomposes brain activity in terms of contributions from multi-scale patterns of coactivation using the network organisation of the structural connectome (Figure 5).

**Figure 5.**
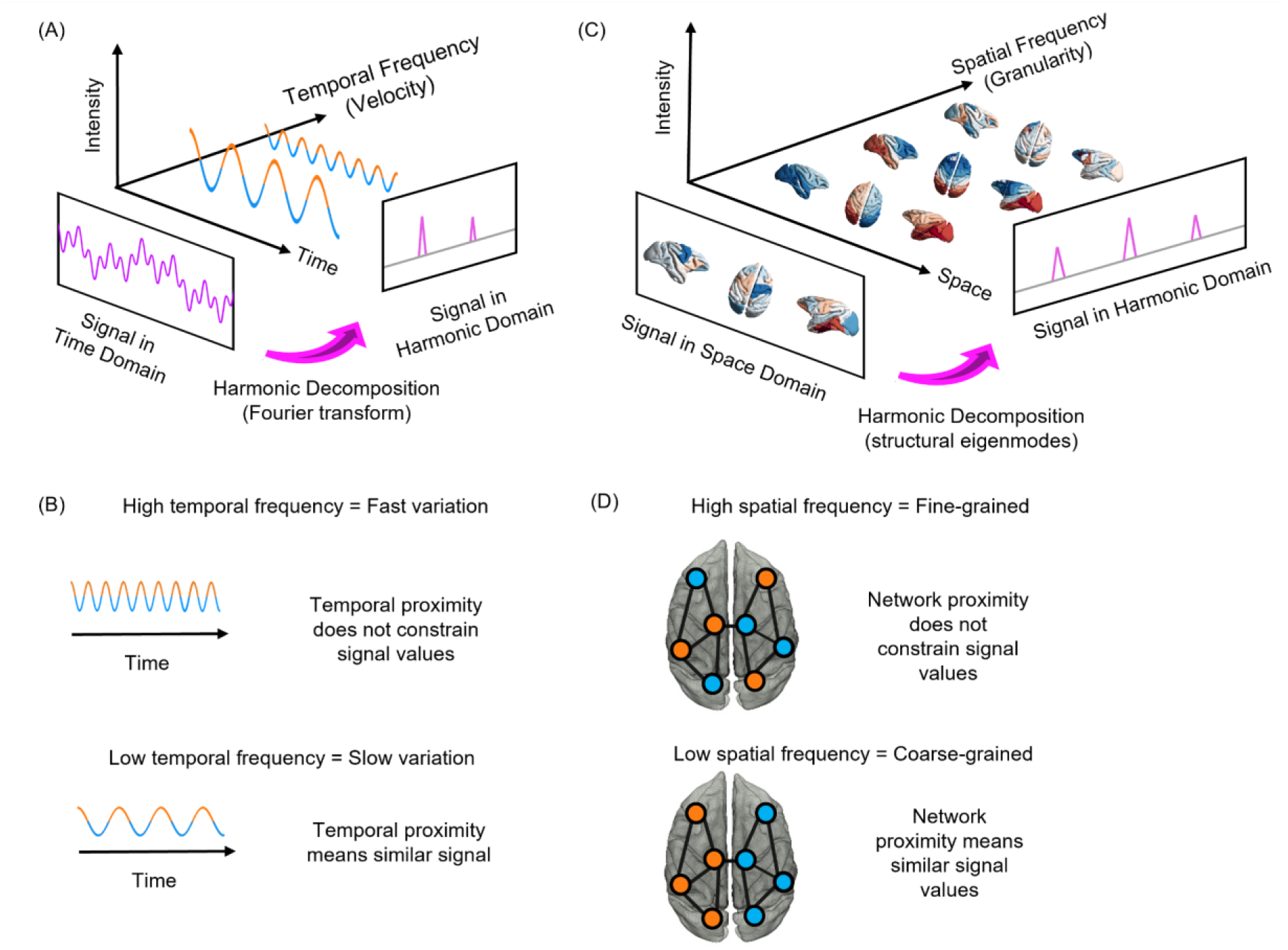
Structural eigenmode decomposition generalises the Fourier transform to the network structure of the brain. (A) In traditional Fourier analysis, a signal in the time domain (represented in terms of successive time points) is decomposed into temporal harmonics of different frequencies and thereby rendered in terms of a new set of basis functions. (B) High-frequency temporal harmonics correspond to rapidly varying signals, such that data points may have very different values even if they are close in time. In contrast, low-frequency temporal harmonics correspond to signals that change slowly over time, such that temporally contiguous have similar values, reflecting a greater time dependence of the signal. (C) Harmonic decomposition of the connectome involves decomposing a signal in the spatial domain (represented in terms of fMRI activation at discrete spatial locations over the cortex) into harmonic modes of the structural connectome, resulting in a new set of basis functions in terms of whole-brain distributed patterns of activity propagation distributed throughout the brain at different spatial scales (granularity), from global patterns of smooth variation along geometrical axes (left–right and anterior– posterior being the most prominent) to increasingly localised patterns. Note that here, frequency is not about time, but about spatial scale. (D) Low-frequency (coarse-grained) connectome harmonics indicate that the spatial organisation of the functional signal closely matches the underlying organisation of the structural connectome: Nodes that are strongly connected exhibit similar functional signals (indicated by colour). High-frequency (fine-grained) patterns indicate divergence between the spatial organisation of the functional signal and the underlying network structure, where nodes may exhibit different functional signals even if they are closely connected in the structural network (Luppi, Vohryzek, *et al*., 2023).

Each harmonic mode is a cortex-spanning activation pattern (eigenmode of the structural connectome) characterised by a specific granularity (spatial frequency, Figure 6): large-scale, coarse-grained patterns (e.g., left and right or front and back of the brain) are strongly constrained by the organisation of the structural connectome, such that strongly interconnected regions are predicted to exhibit similar activation (Luppi, Vohryzek, *et al*., 2023). In contrast, high-frequency harmonic modes are relatively unconstrained by the underlying network structure, such that regions may exhibit different activity even if they are strongly connected in the structural network (Figure 5). Therefore, decomposing functional MRI signals in terms of contributions from different connectome harmonics provides a quantification of the extent to which brain activity is constrained by the network structure of the connectome. In other words, harmonic mode decomposition is analogous to the well-known Fourier decomposition, but operating in the spatial domain (Luppi, Vohryzek, *et al*., 2023).

**Figure 6.**
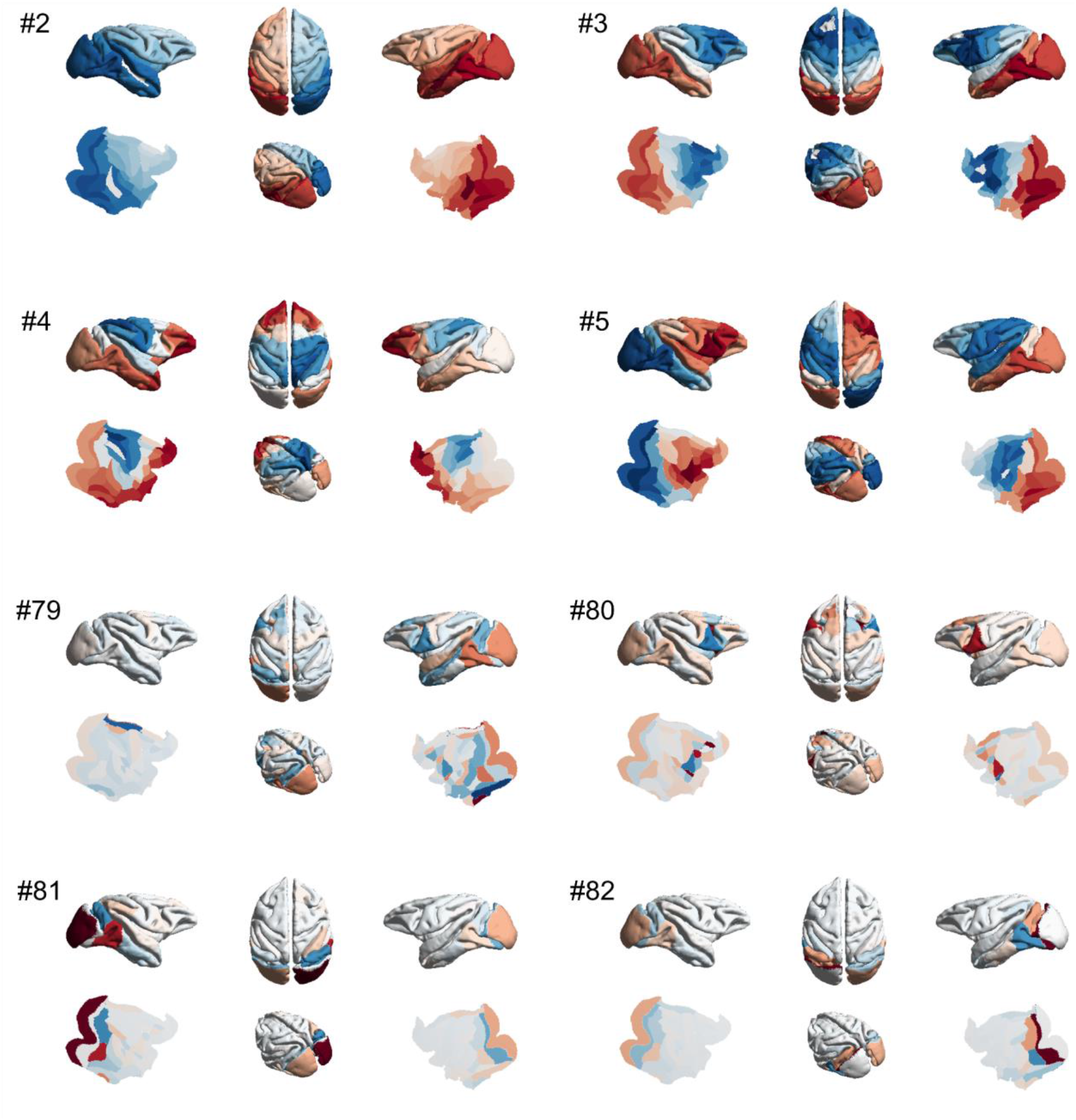
Harmonic modes of the macaque structural connectome. The first four non-uniform harmonic modes of the macaque structural connectome and the last four are shown on the surface of the macaque brain. Note that the first, low-frequency harmonics reveal large-scale patterns, while the last, high-frequency harmonics correspond to localised patterns. The total number of harmonics, and therefore the maximum resolution, corresponds to the number of brain regions in the connectome (here 82). Please note that in the original formulation of Atasoy and colleagues (2016), “connectome harmonics” are specifically defined as the harmonic modes of a high-resolution human structural connectome, obtained from combining long-range white matter tracts and local connectivity within the grey matter. Here we use instead the harmonic modes obtained from a parcellated macaque connectome. To avoid confusion, we refer to the eigenmodes obtained in this way as “harmonic modes”.

The use of connectome-specific harmonic decomposition of cortical activity allows us to quantify the contribution of structural organisation to brain activity, across different spatial scales: from large-scale to localised. This approach therefore goes beyond previous investigations that simply assessed the similarity of structural and functional connectivity at a single scale (Barttfeld et al., 2015; Demertzi et al., 2019; Gutierrez-Barragan et al., 2021). Specifically, for each time-point, the magnitude of contribution of each harmonic to the BOLD activation across the brain (weighted by its associated eigenvalue) is termed the energy (see Methods). Through this formalism, we tested whether the different anaesthetics induced changes in the contribution of structural connectivity to functional activity, and whether such changes (if any) would be reversed upon restoration of responsiveness by thalamic DBS.

Our results indicate that the energy of connectome harmonics in the macaque is reshaped by loss and recovery of consciousness. In particular, the energy of connectome harmonics is significantly increased in the macaque brain under deep (but not light) anaesthesia, whether induced by sevoflurane, propofol, or ketamine (Figure 7 and Figure S5). Remarkably, this pattern is replicated in an independent dataset of propofol anaesthesia, and it is reversed under the effects of thalamic stimulation with both amplitude and site dependence: high amplitude (5V) stimulation of the centro-median thalamic nucleus brought the overall harmonic energy significantly back closer to awake levels, whereas the same stimulation delivered at low amplitude (3V) induced a significant but smaller effect. When the same electrical input (both low and high amplitude) was delivered to the ventro-lateral part of the thalamus, the overall harmony energy did not significantly differ from the anaesthesia condition (Figure 7).

**Figure 7.**
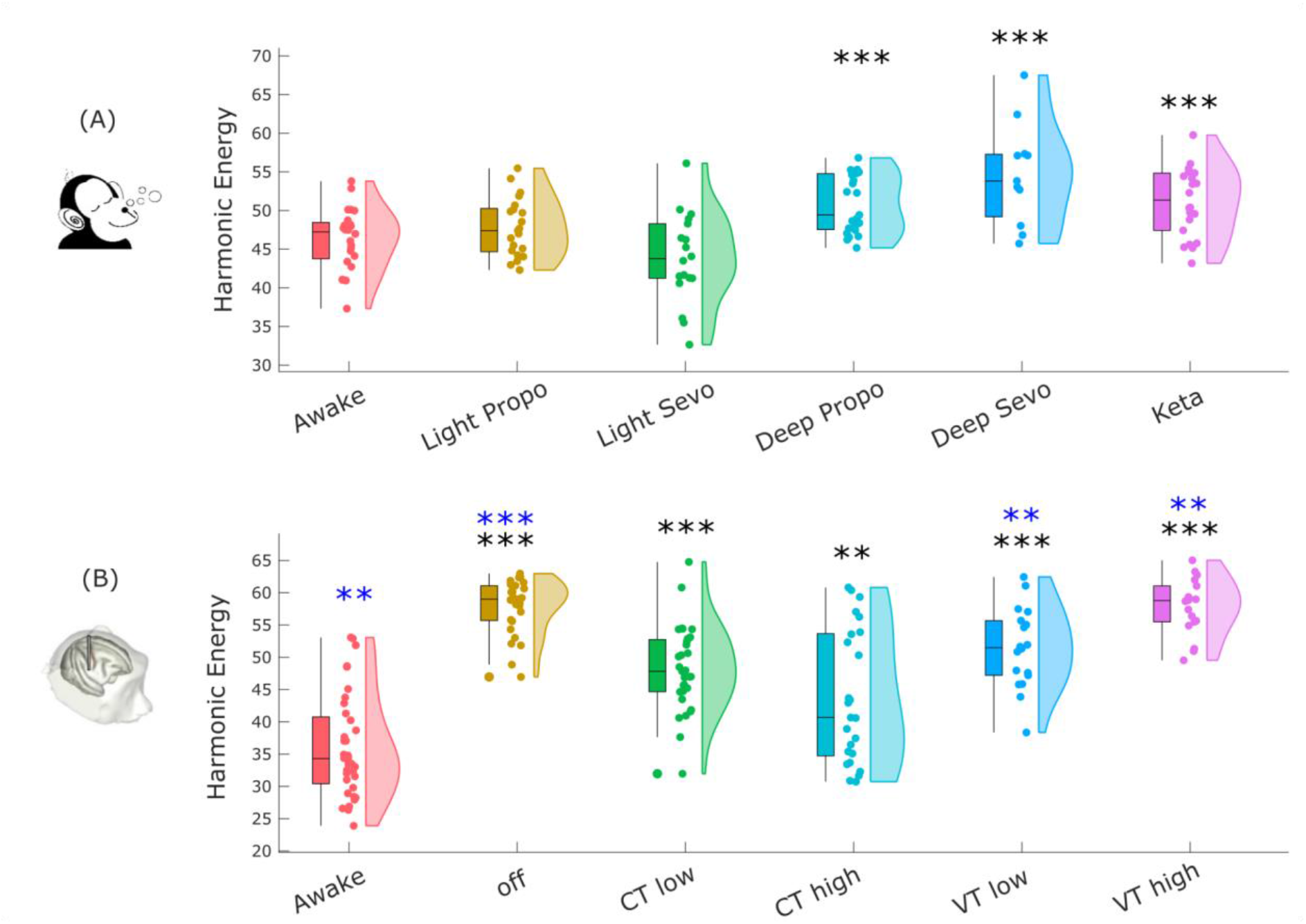
Energy of harmonic modes in the macaque is reshaped by loss and recovery of responsiveness. (A) Energy of connectome harmonics is significantly increased in the macaque brain under deep (but not light) anaesthesia, whether induced by sevoflurane, propofol, or the dissociative anaesthetic ketamine. (B) Effect of thalamic deep-brain stimulation on the energy of connectome harmonics during anaesthesia. Box plots indicate the median and interquartile range of the distribution. * *p* < 0.05; ** *p* < 0.01; *** *p* < 0.001 (FDR-corrected), compared against Awake condition; * *p* < 0.05; ** *p* < 0.01; *** *p* < 0.001 (FDR-corrected), compared against CT high condition; see Figure S5 for comparisons between all pairs.

### Multi-dimensional Representation of Loss and Recovery of Consciousness

Finally, we bring together our three complementary analyses to generate a multi-dimensional profile that might better characterise perturbations of consciousness. We show this by using the magnitude of effect size for each condition of perturbed consciousness compared with wakefulness (Figure 8). For the multi-anaesthetic dataset, we find that hierarchical integration displays the largest effect sizes across all anaesthetics; additionally, we clearly see the greater effect of deep anaesthesia, with all three drugs (propofol, sevoflurane, and ketamine) pushing the effect sizes further away from zero (Figure 8A). Interestingly, hierarchical integration was also strongly affected by high amplitude electrical stimulation of the centro-median thalamus, essentially reducing the effect of anaesthesia back to a near-zero effect size, when compared with wakefulness (Figure 8B). In general, this experimental condition was the most effective in counteracting the effects of propofol on the distributed functional signatures (indicated by the smallest effect sizes when compared against wakefulness), except for the range of the principal FC gradient, which was more restored by low-rather than high-amplitude stimulation of the centro-median thalamic nucleus (Figure 8B). Overall, this multi-dimensional representation in terms of structural and functional eigenmode reorganisation provides a compact description, identifying relevant axes along which perturbations of consciousness can manifest.

**Figure 8.**
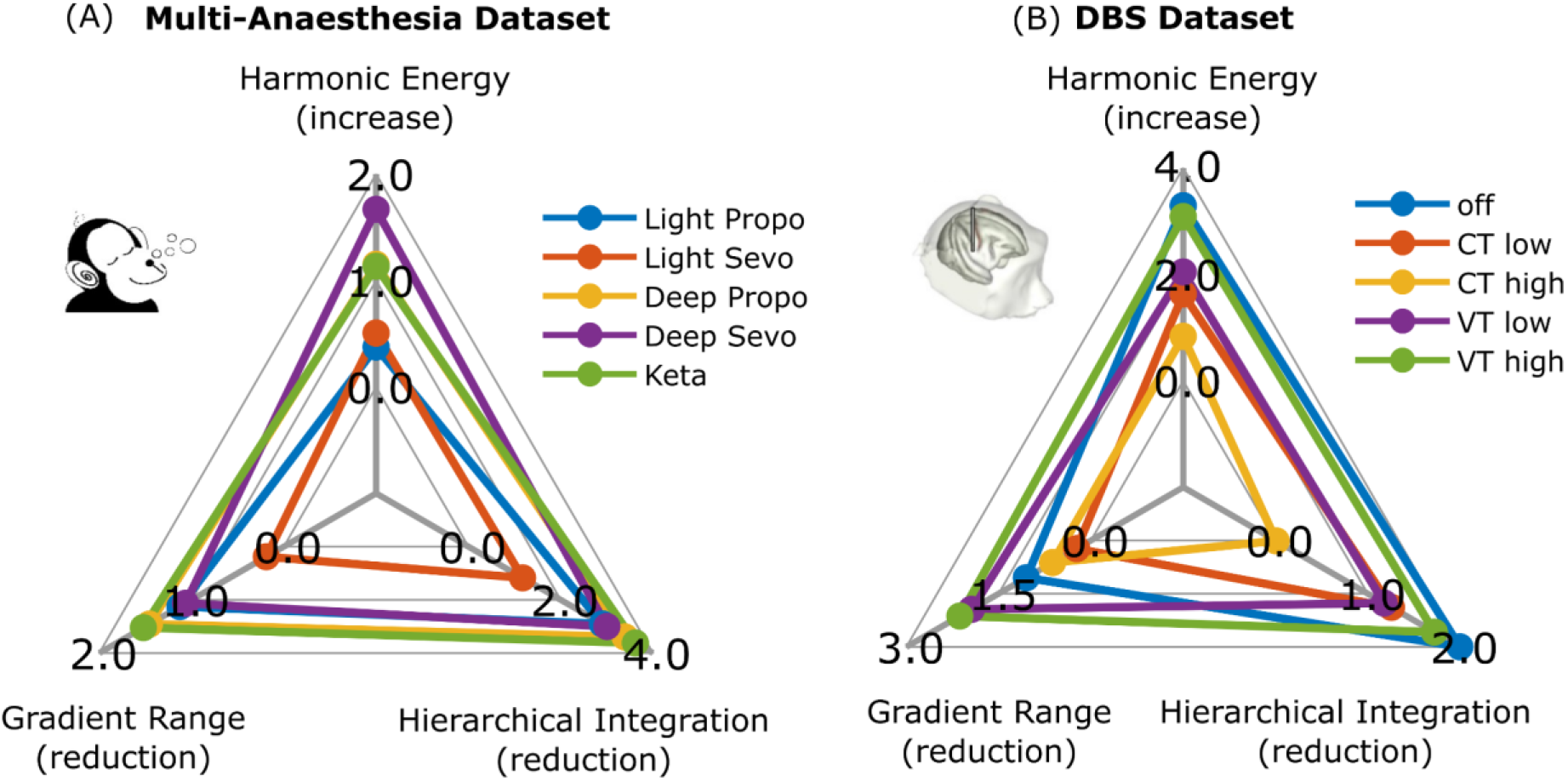
Representing perturbations of consciousness in terms of structural and functional eigenmode reorganisation. (A): Radial plot showing the magnitude (absolute value) of the effect size obtained when comparing each state of perturbed consciousness against wakefulness for the multi-anaesthesia data set. (B) Radial plot shows the magnitude (absolute value) of the effect size obtained when comparing each state of perturbed consciousness against wakefulness, for the DBS dataset.

## Discussion

Here, we capitalised on two unique fMRI datasets of macaque brains during anaesthesia (propofol, ketamine, sevoflurane) with versus without simultaneous intracranial electrical stimulation of different thalamic nuclei. We investigated anaesthetic-induced reorganisation of macaque brain functional architecture, its relationship to brain structure across brain scales, and its restoration by causal intervention (electrical thalamic stimulation). Our goals were twofold. First, we sought to identify consistent signatures of consciousness in terms of distributed brain patterns, across different ways of consciousness loss, by considering anaesthetics with distinct molecular mechanisms. Second, we aimed to determine whether the reorganisation of distributed brain function consistently induced by different anaesthetics can be reversed by targeted stimulation of different subregions of the thalamus.

Thanks to the broad spatial coverage provided by fMRI, we were able to characterise distributed brain function in terms of its relationship to the underlying structural organisation and in terms of how it supports different facets of hierarchical organisation. We showed that large-scale patterns are reliably altered when consciousness is suppressed, regardless of the anaesthetic agent, and that they are restored upon DBS-induced awakening, even in the presence of continued anaesthetic infusion. Taken together, these observations indicate that such changes are neither specific to a particular anaesthetic agent nor exclusively related to the mere presence of the anaesthetic in the bloodstream: rather, they are associated with behavioural markers of wakefulness, with the stimulation results being both location– and amplitude-dependent (Tasserie *et al*., 2022).

We examined how anaesthetic-induced unconsciousness reshapes distributed functional organisation of the primate brain, by studying a dataset of functional MRI recordings from macaque monkeys anaesthetised with propofol, sevoflurane, or ketamine (see the Methods section and (Uhrig *et al*., 2018) for details). Previous work has consistently shown increased correspondence between structural and functional connectivity under anaesthesia (Barttfeld *et al*., 2015; Uhrig *et al*., 2018; Demertzi *et al*., 2019; Gutierrez-Barragan *et al*., 2022; Tasserie *et al*., 2022). However, existing investigations of structure-function correspondence during anaesthesia have typically relied on correlation and analogous distance metrics, which operate at a single spatial scale, whereas it is well established that brain structure and function both exhibit multi-scale, network organisation. This motivates the use of a multi-scale approach, provided by the framework of structural eigenmode decomposition, which simultaneously considers all available scales of organisation: from a single region to the entire cortex). Our results showed that increased multi-scale coupling between brain activity and structural eigenmodes, as recently observed in the human brain during pharmacological and pathological loss of consciousness (Luppi et al., 2023), can be generalised to the macaque brain – thereby supporting previous results that operated at a single scale (and which were obtained in terms of functional connectivity rather than activity).

We also showed that the signatures that we identified generalise across anaesthetic agents, including ketamine. Unlike sevoflurane and propofol, ketamine does not act primarily as an agonist of GABA-A receptors, but rather as an antagonist of NMDA receptors. Note that this result is not in contrast with the ketamine-induced decreased structural coupling observed in humans by Luppi and colleagues (2023), because that study employed a sub-anaesthetic dose of ketamine (i.e., having psychedelic-like effects), whereas the dose of ketamine used in the present study induced general anaesthesia. Thus, showing that previously identified signatures of unconsciousness can be generalised to this additional pharmacological perturbation, improve the robustness of consciousness-specificity of these signatures, while also suggesting that ketamine can have opposite effects on structure-function relationships in the brain, depending on its dosage and the corresponding changes in subjective experience.

In the human brain, considering highly fine-grained harmonic modes of the human connectome in the order of 18,000 surface vertices (Luppi et al., 2023), harmonic mode decomposition has revealed that anaesthesia and disorders of consciousness induce a shift in the relative contribution of different harmonics, increasing the contribution of low-frequency (structurally-constrained) harmonics at the expense of high-frequency (liberal) ones. We note that the resolution of harmonic modes is fundamentally limited by the resolution of the underlying structural connectome: in the present case, 82 regions, each spanning several millimetres of cortex. Therefore, even the most fine-grained eigenmodes from the macaque 82-node connectome correspond to dividing the cortex in fewer than 100 patches – a number that lies firmly in the low-frequency (structurally-constrained) range for the human 18,000– eigenmode decomposition. In other words, when only a resolution of fewer than 100 harmonic eigenmodes is considered, the present results coincide with those reported by Luppi et al (2023): unconsciousness manifests as an increased contribution of structural constraints to cortical functional activation. Future work using higher-resolution anatomical connectivity may enable us to obtain finer-grained insights about the respective roles of low-frequency versus high-frequency structural eigenmodes, such as by using the “structural decoupling index” to quantify their balance (Preti and Van De Ville, 2019).

Decomposition in terms of harmonic modes of the structural connectome characterises distributed brain function in terms of the relationship between functional activation and distributed patterns derived from brain anatomical network organisation, showing how this relationship changes when consciousness is lost. However, the harmonic modes themselves do not change, being based on structural connectivity, which is relatively stable at short timescales and unaffected by anaesthesia. Therefore, we complemented our investigation of distributed structure-function relationships with an additional investigation of distributed patterns arising from functional connectivity, termed functional gradients. This alternative perspective identified diminished hierarchical character of brain function under unconsciousness: both in terms of the processing distance (contraction of the principal functional gradient), and in terms of diminished hierarchical integration across nested functional modules, when considering all functional eigenmodes, rather than just the principal one.

We also note that both approaches that considered all eigenmodes (hierarchical integration and harmonic mode decomposition) exhibited greater effect sizes than consideration of the principal gradient (principal functional eigenmode) alone. Indeed, the measure of gradient range also exhibited a deviation from the behavioural pattern, being more influenced by low– than high-amplitude stimulation of the central thalamus – although we note that both low– and high-amplitude CT stimulation were nevertheless clearly superior to stimulation of the control VT site. As previously reported, low-amplitude stimulation of the central thalamus has weaker effect on behaviour, than high-amplitude stimulation (Tasserie et al., 2022). Thus, gradient range is influenced by CT stimulation even before this stimulation is sufficient to induce restoration of responsiveness, whereas high-amplitude stimulation is more closely aligned with behaviour. In contrast, the structural eigenmode decomposition and hierarchical integration, which consider all eigenmodes (structural and functional, respectively), both displayed the largest effect for high-amplitude stimulation of the central thalamus, coinciding with the fullest restoration of consciousness. These observations reinforce the value of considering distributed brain function across multiple scales: the full effects of high-amplitude stimulation appear to be spread across multiple functional eigenmodes, such that only considering the first one provides an incomplete picture with weaker link to behaviour. On the other hand, although here we did not focus on the specific topography of functional gradients, but rather on their extent and relationships, we expect that future work will also benefit from such an approach. Overall, these convergent results from distributed function extend previous observations from macaques and humans under anaesthesia and disorders of consciousness (Signorelli *et al*., 2021; Luppi *et al*., 2022).

Crucially, the restoration of such distributed cortical patterns can be triggered by selective stimulation of a specific subcortical region, as demonstrated by our spatially specific causal intervention: the centro-median thalamic nucleus – in contrast to the much weaker effects elicited by control site stimulation of the ventral lateral nucleus of the thalamus. Therefore, both wakefulness and its associated large-scale cortical functional patterns are under causal control by a very selective locus. Thus, our unique combination of broad recording coverage (from fMRI) and specific causal intervention (from DBS) enables us to show that global functional patterns are orchestrated locally, showing how the locationist and distributed approaches to the neural correlates of consciousness are not antithetical, but rather complement each other. In this sense, our results help reconcile the traditional locationist approach to the neural correlates of consciousness with recent advances in understanding brain function in terms of distributed patterns (Suzuki and Larkum, 2020).

Our findings regarding thalamic control of distributed cortical functions are consistent with recent views on the thalamus as a controller of brain dynamics (Liu *et al*., 2015; Shine, 2019, 2021), and more broadly on its role in supporting consciousness (Lioudyno *et al*., 2013; Vijayan *et al*., 2013; Akeju *et al*., 2014; Flores *et al*., 2017), including connectivity with cortical regions displaying consciousness-specific signatures (Luppi *et al*., 2019) and the effects of selective thalamic stimulation on consciousness and arousal in animals (Alkire *et al*., 2007, 2009; Lewis *et al*., 2015; Bastos *et al*., 2021; Tasserie *et al*., 2022) and human patients (Tononi, 2004; Schiff *et al*., 2007; Staunton, 2008; Mashour *et al*., 2020). Indeed, recent studies also reported that electrical stimulation of specific central-lateral thalamic nuclei (but not control sites) can restore arousal during anaesthetic-induced loss of responsiveness in macaques (Tononi, 2004; Schiff *et al*., 2007; Staunton, 2008; Mhuircheartaigh *et al*., 2010; Lioudyno *et al*., 2013; Ní Mhuircheartaigh *et al*., 2013; Vijayan *et al*., 2013; Akeju *et al*., 2014; Flores *et al*., 2017; Hemmings *et al*., 2019; Kelz *et al*., 2019; Mashour *et al*., 2020; Redinbaugh *et al*., 2020, 2022; Afrasiabi *et al*., 2021; Bastos *et al*., 2021; Gammel *et al*., 2023; Kantonen *et al*., 2023), counteracting the loss of high-frequency (gamma-band) activity and communication between the thalamus and deep cortical layers, which is induced by anaesthesia. However, these studies were limited by the use of electrodes for recording, given the authors’ interest in different frequency bands of neural activity – consequently limiting spatial coverage to 2-8 sites. In contrast, the present work combined thalamic stimulation with global coverage of the entire cortex, thanks to the use of functional MRI. This unique set-up enabled us to study both local and distributed neural contributions to consciousness. Our evidence that selective thalamic DBS can restore not only arousal, but also distributed neural signatures of consciousness that are disrupted across anaesthetics, lends further support for the translational potential of DBS as an avenue for treating the challenging condition of patients suffering from disorders of consciousness (Cohadon and Richer, 1993; Schiff *et al*., 2007; Lemaire *et al*., 2018; Edlow *et al*., 2021) – although with the clear caveat that such patients typically exhibit widespread cortical and subcortical damage, unlike the non-human primates in the present study.

An important limitation of this work is that our distributed approach to brain function does not provide insights about the role of specific brain regions: instead, we focused on different whole-brain patterns. A substantial body of previous work has focused on specific regions, and our goal here was to provide an alternative perspective. Additionally, thanks to the DBS dataset we were still able to identify localised contributions to consciousness. Nevertheless, future analytic approaches capable of simultaneously identifying both local and distributed signatures may hold additional promise for unveiling the rich interplays that support consciousness. Another limitation is that our analysis was exclusively cortical (except for the thalamic stimulation), yet other subcortical structures such as the basal ganglia and brainstem (Moruzzi and Magoun, 1949; Brown, Lydic and Schiff, 2010; Mhuircheartaigh *et al*., 2010; Braun *et al*., 2011; Hudetz, 2012; Hemmings *et al*., 2019; Guang *et al*., 2021; Spindler *et al*., 2021) are known to play an important role in anaesthetic-induced and pathological loss of responsiveness (Bastos *et al*., 2021; Tasserie *et al*., 2022). Future work including subcortical regions may therefore provide additional insights.

Future directions can also combine methods from eigenmode analysis and combinatorial topology to construct efficient representations of high-order information-theoretic signals (Medina-Mardones *et al*., 2021), which are a hallmark of many complex systems including the brain in its different states of consciousness. Additionally, here we relied on a symmetrised version of the macaque connectome, which is required for eigendecomposition in order to ensure real eigenmodes. However, since a directed macaque connectome is available, we expect that generalisation of harmonic mode analysis to account for directed connections, may provide a more refined picture. Pertaining to alternative avenues of eigenmode decomposition, recent work suggested that structural eigenmodes obtained from the geometry of the human brain may outperform those obtained from the human connectome, in terms of explaining variance in the functional signals (Pang *et al*., 2023). Thus, future work may investigate whether cortical geometry provides insights into the reorganisation of consciousness, beyond those provided by the connectome.

In the absence of subjective reports from animals, behavioural markers alone play a more prominent role, although it is known from research in humans that on occasion behaviour and subjective experience can be dissociated – including with ketamine. More broadly, our analysis is based on fMRI acquired in resting state, i.e. in the absence of a stimulus or a task. Though beyond the scope of the present work, analysis of distributed patterns pertaining to the dysfunctional spread of naturalistic and synthetic stimuli, which was also observed in the same anaesthetised animals, will provide a more comprehensive understanding of how such patterns underpin information-processing and its perturbation by anaesthetics. Conversely, a strength of this work is the replication of our results not only across multiple anaesthetics, but also in the independent DBS-fMRI dataset, showing their robustness to acquisition differences such as the presence or absence of MION contrast agent. Likewise, the DBS data enabled us to assess the respective roles of both location (central versus ventral-lateral thalamus) and stimulation strength.

Overall, we showed that distributed patterns of functional activity and connectivity of the primate brain provide rich characterisations of consciousness, its suppression by different anaesthetics, and its restoration by thalamic stimulation. The resulting insights align both with the well-known increase in structure-function coupling observed across a variety of pharmacological and pathological states of unconsciousness across species, and also with prominent theories of consciousness that postulate a central role for the integration of information, which we show here to be reliably disrupted in the unconscious primate brain. We have also provided two convergent lines of evidence for a reduced hierarchical character of cortical functions in the unconscious primate brain.

## Materials and Methods

### Animals

For the anaesthesia dataset, the details of the acquisitions have been previously reported in (Barttfeld *et al*., 2015; Uhrig *et al*., 2018; Signorelli *et al*., 2021). Five rhesus macaques were included for analyses (Macaca mulatta, one male, monkey J, and four females, monkey A, K, Ki, and R, 5-8 kg, 8-12 yr of age), in a total of six different arousal conditions: awake state, ketamine, light propofol, deep propofol, light sevoflurane, and deep sevoflurane anaesthesia. Three monkeys were used for each condition: awake state (monkeys A, K, and J), ketamine (monkeys K, R and Ki), propofol (monkeys K, R, and J), sevoflurane (monkeys Ki, R, and J). Each monkey had fMRI resting-state acquisitions on different days and several monkeys were scanned in more than one experimental condition. All procedures are in agreement with the European Convention for the Protection of Vertebrate Animals used for Experimental and Other Scientific Purposes (Directive 2010/63/EU) and the National Institutes of Health’s Guide for the Care and Use of Laboratory Animals. Animal studies were approved by the institutional Ethical Committee (Commissariat à l’Energie atomique et aux Énergies alternatives; Fontenay aux Roses, France; protocols CETEA \#10-003 and 12-086).

For the DBS dataset, details were provided in (Tasserie *et al*., 2022). Five male rhesus macaques (Macaca mulatta, 9 to 17 years and 7.5 to 9.1 kg) were included, three for the awake (non-DBS) experiments (monkeys B, J, and Y) and two for the DBS experiments (monkeys N and T). All procedures are in agreement with 2010/63/UE, 86-406, 12-086 and 16-040.

### Anaesthesia Protocol

The anaesthesia protocol is thoroughly described in previous studies (Barttfeld *et al*., 2015; Uhrig *et al*., 2018). Monkeys received anesthesia either with ketamine (Uhrig *et al*., 2018), propofol (Barttfeld *et al*., 2015; Uhrig *et al*., 2018) or sevoflurane (Barttfeld *et al*., 2015; Uhrig *et al*., 2018), with two different levels of anesthesia for propofol and sevoflurane anesthesia (light and deep). The anesthesia levels were defined according to the monkey sedation scale, based on spontaneous movements and the response to external stimuli (presentation, shaking or prodding, toe pinch), and corneal reflex (Uhrig, 2016). For each scanning session, the clinical score was determined at the beginning and end of the scanning session, together with continuous electroencephalography monitoring (Uhrig *et al*., 2016). Monkeys were intubated and ventilated as previously described (Barttfeld *et al*., 2015; Uhrig *et al*., 2018). Heart rate, noninvasive blood pressure, oxygen saturation, respiratory rate, end-tidal carbon dioxide, and cutaneous temperature were monitored (Maglife, Schiller, France) and recorded online (Schiller).

During ketamine, deep propofol, and deep sevoflurane anesthesia, monkeys stopped responding to all stimuli, reaching a state of general anesthesia. For ketamine anesthesia, ketamine was injected intramuscular (20 mg/kg; Virbac, France) for induction of anesthesia, followed by a continuous intravenous infusion of ketamine (15 to 16 mg · kg–1 · h–1) to maintain anesthesia. Atropine (0.02 mg/kg intramuscularly; Aguettant, France) was injected 10 min before induction, to reduce salivary and bronchial secretions. For propofol anesthesia, monkeys were trained to be injected an intravenous propofol bolus (5 to 7.5 mg/kg; Fresenius Kabi, France), followed by a target-controlled infusion (Alaris PK Syringe pump, CareFusion, USA) of propofol (light propofol sedation, 3.7 to 4.0 µg/ml; deep propofol anesthesia, 5.6 to 7.2 µg/ml) based on the “Paedfusor” pharmacokinetic model (Absalom and Kenny, 2005). During sevoflurane anesthesia, monkeys received first an intramuscular injection of ketamine (20 mg/kg; Virbac) for induction, followed by sevoflurane anesthesia (light sevoflurane, sevoflurane inspiratory/expiratory, 2.2/2.1 volume percent; deep sevoflurane, sevoflurane inspiratory/expiratory, 4.4/4.0 volume percent; Abbott, France). Only 80 minutes after the induction, the scanning sessions started for the sevoflurane acquisitions to get a washout of the initial ketamine injection (Schroeder *et al*., 2016). To avoid artefacts related to potential movements throughout magnetic resonance imaging acquisition, a muscle-blocking agent was coadministered (cisatracurium, 0.15 mg/kg bolus intravenously, followed by continuous intravenous infusion at a rate of 0.18 mg · kg–1 · h–1; GlaxoSmithKline, France) during the ketamine and light propofol sessions.

For the DBS dataset, anaesthesia was induced with an intramuscular injection of ketamine (10 mg/kg; Virbac, France) and dexmedetomidine (20 μg/kg; Ovion Pharma, USA) and then the same method as described above for deep propofol sedation was used (Monkey T: TCI, 4.6 to 4.8 µg/ml; monkey N: TCI, 4.0 to 4.2 µg/ml). Two monkeys were implanted with a clinical DBS electrode (Medtronic, Minneapolis, MN, USA, lead model 3389).The DBS protocol is thoroughly described in Tasserie et al, 2022. The right centro-median thalamus was targeted by stereotactic surgery using a neuro-navigation system (BrainSight, Rogue, Canada), guided by the rhesus macaque atlases (paxinos et al., Academic press 2008 and Saleem Logothetis Academic Press 2012) and a preoperative and intraoperative anatomical MRI.

### Functional Magnetic Resonance Imaging Data Acquisition

For the awake condition, monkeys were implanted with a magnetic resonance compatible head post and trained to sit in the sphinx position in a primate chair (Uhrig, Dehaene and Jarraya, 2014)). For the awake scanning sessions, monkeys sat inside the dark magnetic resonance imaging scanner without any task and the eye position was monitored at 120 Hz (Iscan Inc., USA). The eye-tracking was performed to make sure that the monkeys were awake during the whole scanning session and not sleeping. The eye movements were not regressed out from rfMRI data. For the anesthesia sessions, animals were positioned in a sphinx position, mechanically ventilated, and their physiologic parameters were monitored. No eye-tracking was performed in anesthetic conditions. For the anesthesia dataset, before each scanning session, a contrast agent, monocrystalline iron oxide nanoparticle (Feraheme, AMAG Pharmaceuticals, USA; 10 mg/kg, intravenous), was injected into the monkey’s saphenous vein (Vanduffel *et al*., 2001). Monkeys were scanned at rest on a 3-Tesla horizontal scanner (Siemens Tim Trio, Germany) with a single transmit-receive surface coil customized to monkeys. Each functional scan consisted of gradient-echo planar whole-brain images (repetition time = 2,400 ms; echo time = 20 ms; 1.5-mm3 voxel size; 500 brain volumes per run).

For the DBS dataset, monkeys were scanned at rest on a 3-Tesla horizontal scanner (Siemens, Prisma Fit, Erlanger Germany) with a customized eight-channel phased– array surface coil (KU Leuven, Belgium). The parameters of the functional MRI sequences were: echo planar imaging (EPI), TR = 1250 ms, echo time (TE) = 14.20 ms, 1.25-mm isotropic voxel size and 500 brain volumes per run.

### Data preprocessing and time series extraction

For the anaesthesia dataset, a total of 157 functional magnetic imaging runs were acquired (Uhrig *et al*., 2018): Awake, 31 runs (monkey A, 4 runs; monkey J, 18 runs; monkey K, 9 runs), Ketamine, 25 runs (monkey K, 8 runs; monkey Ki, 7 runs; monkey R, 10 runs), Light Propofol, 25 runs (monkey J, 2 runs; monkey K, 10 runs; monkey R, 12 runs), Deep Propofol, 31 runs (monkey J, 9 runs; monkey K, 10 runs; monkey R, 12 runs), Light Sevoflurane, 25 runs (monkey J, 5 runs; monkey Ki, 10 runs; monkey R, 10 runs), Deep Sevoflurane anaesthesia, 20 runs (monkey J, 2 runs; monkey Ki, 8 runs; monkey R, 11 runs). For details, check the supplementary tables for (Barttfeld *et al*., 2015; Uhrig *et al*., 2018; Signorelli *et al*., 2021) (http://links.ww.com/ALN/B756).

Functional images were reoriented, realigned, and rigidly coregistered to the anatomical template of the monkey Montreal Neurologic Institute (Montreal, Canada) space with the use of Python programming language and Oxford Centre Functional Magnetic Resonance Imaging of the Brain Software Library software (United Kingdom, http://www.fmrib.ox.ac.uk/fsl/; accessed February 4, 2018) (Uhrig, Dehaene and Jarraya, 2014)). From the images, the global signal was regressed out to remove any confounding effect due to physiologic changes (e.g., respiratory or cardiac changes).

For the DBS dataset, a total of 199 Resting State functional MRI runs were acquired: Awake 47 runs (monkey B: 18 runs; monkey J: 13 runs; monkey Y: 16 runs), anaesthesia (DBS-off) 38 runs (monkey N: 16 runs,; monkey T: 22 runs), low amplitude centro-median thalamic DBS 36 runs (monkey N: 18 runs; monkey T: 18 runs), low amplitude ventro-lateral thalamic DBS 20 runs (monkey T), high amplitude centro-median thalamic DBS 38 runs (monkey N: 17 runs; monkey T: 21 runs), and high amplitude ventro-lateral thalamic DBS 20 runs (monkey T: 20 runs).

Images were preprocessed using Pypreclin (Python preclinical pipeline) (Tasserie et al., Neuroimage 2020). Functional images were corrected for slice timing and B0 inhomogeneities, reoriented, realigned, resampled (1.0 mm isotropic), masked, coregistered to the MNI macaque brain template (Frey *et al*., 2011), and smoothed (3.0-mm Gaussian kernel). Anatomical images were corrected for B1 inhomogeneities, normalised to the anatomical MNI macaque brain template, and masked.

Data were parcellated according to the Regional Map parcellation (Kötter and Wanke, 2005). This parcellation comprises 82 cortical ROIs (41 per hemisphere; Supplementary Table 4). Voxel time series were filtered with low-pass (0.05-Hz cutoff) and high-pass (0.0025-Hz cutoff) filters and a zero-phase fast-Fourier notch filter (0.03 Hz) to remove an artifactual pure frequency present in all the data (Barttfeld *et al*., 2015; Uhrig *et al*., 2018).

Furthermore, an extra cleaning procedure was performed to ensure the quality of the data after time-series extraction (Signorelli *et al*., 2021). The procedure was based on a visual inspection of the time series for all the nodes, the Fourier transform of each signal, the functional connectivity for each subject, and the dynamical connectivity computed with phase correlation. Trials were kept when the row signal did not present signs of artifactual activity, functional connectivity was coherent with the average and dynamical connectivity presented consistent patterns across time.

Finally, for the anaesthesia data set a total of 119 runs are analysed in subsequent sections: awake state 24 runs, ketamine anaesthesia 22 runs, light propofol anaesthesia 21 runs, deep propofol anaesthesia 23 runs, light sevoflurane anaesthesia 18 runs, deep sevoflurane anaesthesia 11 runs. For the DBS data set, a total of 156 runs are analysed in subsequent sections: awake state 36 runs, Off condition 28 runs, On 3V 31 runs, On 3V control 18 runs, On 5V 25 runs, On 5V control 18 runs.

### Anatomical Parcellation and Structural Connectivity

Anatomical (structural) connectivity data were derived from the recent macaque connectome of (Shen *et al*., 2019), which combines diffusion MRI tractrography with axonal tract-tracing studies, representing the most complete representation of the macaque connectome available to date.

Structural (i.e., anatomical) connectivity data are expressed as a matrix in which the 82 cortical regions of interest are displayed in x-axis and y-axis. Each cell of the matrix represents the strength of the anatomical connection between any pair of cortical areas.

### Structure-Function Coupling via Harmonic Mode Decomposition

Functional spatiotemporal patterns of neural activity derived from fMRI can be decomposed in terms of anatomically-based distributed building blocks: eigenvectors of the graph Laplacian of the structural connectome (Atasoy *et al*., 2017; Atasoy, Deco, *et al*., 2018; Atasoy, Vohryzek, *et al*., 2018). Here we are inspired by the original Connectome Harmonic Decomposition formulated by of Atasoy and colleagues (Atasoy, Donnelly and Pearson, 2016), who used the harmonic modes of a high-resolution human structural connectome, obtained from combining long-range white matter tracts and local connectivity within the grey matter. Since here we use instead the harmonic modes obtained from a parcellated macaque connectome of diffusion MRI and tract-tracing, we refer to the eigenmodes obtained in this way as “harmonic modes”, and reserve the term “connectome harmonics” for those developed by Atasoy and colleagues.

Following the method developed by Atasoy and colleagues (Atasoy, Donnelly and Pearson, 2016), we compute the symmetric graph Laplacian Δ*_G_* on the matrix *C* that represents the macaque structural connectome. To ensure symmetry of the corresponding connectivity matrix, and real eigenvalues, we averaged entries above and below the diagonal to obtain an undirected connectome. Thereafter, we estimate the connectome Laplacian (discrete counterpart of the Laplace operator Δ applied to the network of the macaque structural brain connectivity):

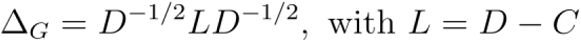

where *D* is the diagonal “degree matrix” of the graph *C* i.e.

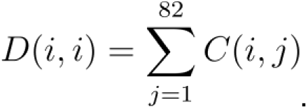

We then calculate the harmonic modes 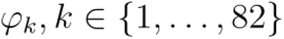 by solving the following eigenvalue equation:

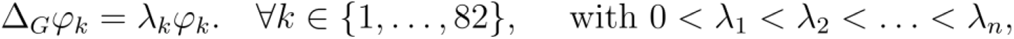

where 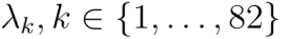 is the corresponding eigenvalue of the eigenvector 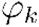. In other words, 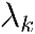 and 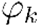 are the eigenvalues and eigenvectors of the Laplacian of the primate structural connectivity matrix (macaque connectome). Therefore, if 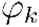 is the harmonic pattern of the 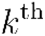 spatial frequency, then the corresponding eigenvalue 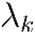 is a term relating to the intrinsic energy of that particular harmonic mode (see figure 1). With an increasing harmonic number 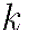, we obtain more complex and fine-grained spatial patterns (see Figures 5 and 6).

### Decomposition of fMRI data

At each timepoint 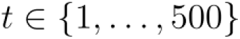 (corresponding to one TR), the spatial pattern of cortical activity over brain regions at time *t*, denoted as *F_t_*, was decomposed as a linear combination of the set of harmonic modes 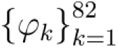:

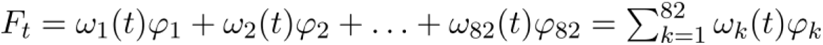

with the contribution 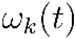 of each harmonic mode 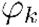 at time *t* being estimated as the projection (dot product) of the fMRI data *F_t_* onto 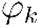:

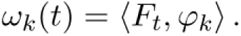

### Energy of harmonic modes

Once the fMRI cortical activation pattern at time has been decomposed into a linear combination of harmonic modes, the magnitude of each harmonic’s contribution to the cortical activity of each harmonic 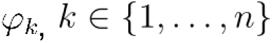, (regardless of sign) at any given timepoint *t*, denoted 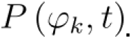, is called its “power”, for analogy with the Fourier transform, is computed as the amplitude of its contribution:

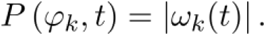

The normalized frequency-specific contribution of each harmonic 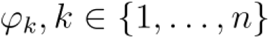 at timepoint *t*, termed “energy”, is estimated by combining the magnitude strength of activation (power) of a particular harmonic mode with its own intrinsic energy given by the associated eigenvalue 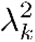:

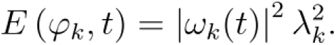

### Functional Gradient Mapping

Macaque cortical functional gradients were calculated using the BrainSpace toolbox https://github.com/ MICA-MNI /BrainSpace (Vos de Wael *et al*., 2020)as implemented in MATLAB, with default parameters for kernel, similarity metric, and sparsity (see below).

We calculate the functional connectivity matrix (FC) as the Pearson correlation between each pair of regional fMRI signals per scan per condition.

Following previous work, each matrix was z-transformed and thresholded row-wise to achieve 90% sparsity, retaining only the strongest connections in each row (Girn *et al*., 2022). The cosine similarity matrix was calculated on the thresholded z-matrix to generate a similarity matrix reflecting the similarity in whole-brain connectivity patterns between vertices. While the FC matrix reflects how similar each pair of regions are in terms of their temporal cofluctuations, this similarity matrix reflects how similar two regions are in terms of their patterns of FC. The similarity matrix is required as input to the diffusion map embedding algorithm that we have used here in agreement with previous work on functional gradients and how they are reshaped by pharmacological interventions (Girn *et al*., 2022).

Diffusion Map Embedding is a nonlinear manifold learning technique (Coifman *et al*., 2005; Margulies *et al*., 2016), that exploits the properties of the graph Laplacian to model the diffusion process, and is therefore related to harmonic mode decomposition – though performed on functional data rather than using a common structural connectome, to reveal its contributions to the functional activations (indeed, there is a deep mathematical analogy exists between diffusion graph embedding on the functional connectome and the recent extension of CHD called “functional harmonics” (Glomb *et al*., 2021; Lioi *et al*., 2021).

The relative influence of density of sampling points on the manifold is controlled by an additional parameter in the range of 0 to 1, which for diffusion map embedding is set to 0.5 to provide a balance between local and global contributions to the embedding space estimation (Coifman *et al*., 2005).

The high-dimensional similarity matrix is treated as a graph, with “connections” (entries of the similarity matrix) reflecting the similarity between the regional patterns of FC. The technique estimates a low-dimensional set of embedding components (gradients); in this low–dimensional space, proximity reflects similarity of the patterns of FC: regions with similar FC patterns (which are strongly connected in the network) are placed close to each other, and regions with low similarity are placed far apart. In this way, each gradient represents one dimension of covariance in the inter-regional similarity between FC patterns, with a small number of gradients capturing most of the dimensions of inter-regional similarity, which can then be visualised by a low-dimensional scatter plot (Figure 4) (Coifman *et al*., 2005; Margulies *et al*., 2016). In the embedding space, each gradient can be understood to be “anchored” at regions that have the strongest values for that gradient, suggesting that this particular embedding dimension captures their similarity profiles of FC well. In contrast, regions that are close to the origin (i.e. have a low absolute value for a particular gradient) mean that they are only minimally similar to the “anchor points” of that gradient, which overall does not strongly capture their FC similarity profile well overall (Coifman *et al*., 2005; Margulies *et al*., 2016).

Therefore, the more different the extremes of a gradient differ along the axis of the gradient, the more the differentiation between regions is being captured by that gradient. To quantify this formally, we calculated the difference between the maximum and minimum values of each scan along the first gradient (Girn *et al*., 2022) (which mathematically captures most of the variation in FC profiles within each scan) and compared these differences across conditions. Note that the dimension of greatest variability (first gradient) is assessed in a data-driven manner and need not be identical across different scans.

### Hierarchical Integration

A third perspective on distributed brain function that can be obtained from studying the brain’s eigenmodes emerges from its hierarchical organisation, and how the latter supports the balance between integration and segregation across scales – which is a central feature of many prominent scientific theories of consciousness.

Classical graph-theoretic measures such as small-worldness and modularity quantify integration and segregation at a single scale and are not suitable for capturing these properties across multiple hierarchical modules (Newman, 2006; Rubinov and Sporns, 2010). However, a recently introduced formalism based on eigenmodes of functional connectivity can provide such quantification (Wang *et al*., 2021).

The functional connectivity (FC) is a symmetric matrix, which can be decomposed as *FC* = *U*Ʌ*U^T^*, where *U* is an orthogonal matrix whose columns are eigenvectors (eigenmodes) of FC, and Ʌ is a diagonal matrix whose entries are the eigenvalues of FC. Each eigenmode of functional connectivity identifies a distinct pattern of regions that are jointly activated (same sign) or alternate (opposite sign). Therefore, hierarchical modules can be identified based on the concordance or discordance of signs between regions across eigenmodes, progressively partitioning the FC into a larger number of modules and submodules, up to the level where each module coincides with a single region, indicative of completely segregated activity. The first eigenmode has the same sign throughout the entire cortex, reflecting global integration. At the next level, two partitions can be detected based on their different signs in the second eigenmode, and each in turn is subdivided at the following level of the hierarchy (i.e., from the third eigenmode) on the basis of regional signs. Thus, segregated modules at one level of the hierarchy can become integrated by being part of the same superordinate module (note that this hierarchical modularity based on eigenmodes is not equivalent to the clustering or modularity maximisation methods (Newman, 2006; Rubinov and Sporns, 2010). During this nested partitioning process, we obtain the module number *M_i_*, (*i* = 1 … *N*) and the modular size *m_j_*, (*i* = 1 . . *M_i_*) at each level.

Each level \textit{i} of the hierarchy is characterised by two quantities: the number of modules *M_i_* into which the cortex is divided, and the covariance explained by the corresponding eigenmode (given by its squared eigenvalue 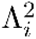). However, the number of modules alone may not properly describe the picture of nested segregation and integration because the size of modules may be heterogeneous. The correction factor was calculated as 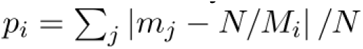 which reflects the deviation from the optimized modular size in the ith level. Thus, the correction effect is stronger for a larger deviation of modular size from homogeneity (Wang *et al*., 2021).

Since the first eigenmode encompasses the entire cortex into one global module, the corresponding eigenvalue 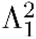 quantifies the overall contribution of global integration, 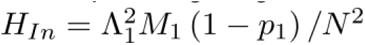. The overall level of segregation across the hierarchy is quantified as 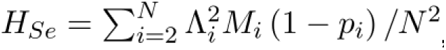, the sum of contributions from all eigenmodes except the first (i.e., all eigenmodes that involve a partitioning of the cortex), each weighted by the corresponding number of modules *M_i_* (further corrected for heterogeneous modular sizes, since a partition into two modules of size 1 and N-1 is clearly not as segregated as a partition into two equally-sized modules).

## Acknowledgments

AIL was supported by a Gates Cambridge Scholarship (OPP 1144), a Travel Grant of the Boehringer Ingelheim Fonds, and the visitor program of the European Institute for Theoretical Neuroscience (EITN). JT was supported by the Fondation pour la Recherche Médicale (FRM grant number ECO20160736100). CMS was supported by FNRS, project MIS/VA – F.4512.21 and grant Embodied-Time – 40011405, Belgium. EAS is supported by the Stephen Erskine Fellowship of Queens’ College, Cambridge. AD and RC are supported by the CNRS and the European Community (Human Brain Project, H2020-945539). BJ was supported by the Fondation Bettencourt Schueller, Fondation de France, Human Brain Project (Corticity project), Institut National de la Santé et de la Recherche Médicale, UVSQ, Commissariat à l’Energie Atomique, Collège de France.

## Author contributions

A.I.L., R.C., A.D., conceived the analysis; J.T., L.U., and B.J. designed the experiments; J.T., L.U., and B.J. collected the data; A.I.L. and C.M.S. analysed the data; A.I.L., C.M.S., and R.C. wrote the manuscript with feedback from all co-authors.. B.J., A.D., E.A.S., supervised the project. All authors approved the manuscript

