## Supplementary Material for "Local Orchestration of Global Functional Patterns Supporting Loss and Restoration of Consciousness in the Primate Brain"

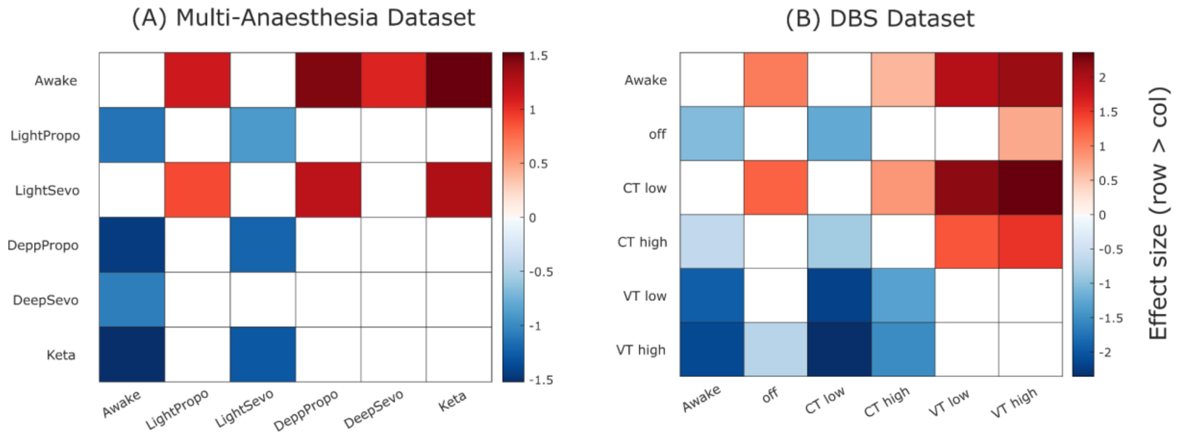


**Figure S1. Effect sizes for significant changes in functional gradient range across states of consciousness**. (A) Matrix shows the effect sizes for each statistical contrast between conditions of the multi-anaesthesia dataset; positive (negative) values indicate that the condition in the row is greater (lower) than the condition in the column. White entries indicate that the difference is not statistically significant. (B) Matrix shows the effect sizes for each statistical contrast between conditions in the DBS dataset; positive (negative) values indicate that the condition in the row is greater (lower) than the condition in the column. White entries indicate that the difference is not statistically significant.

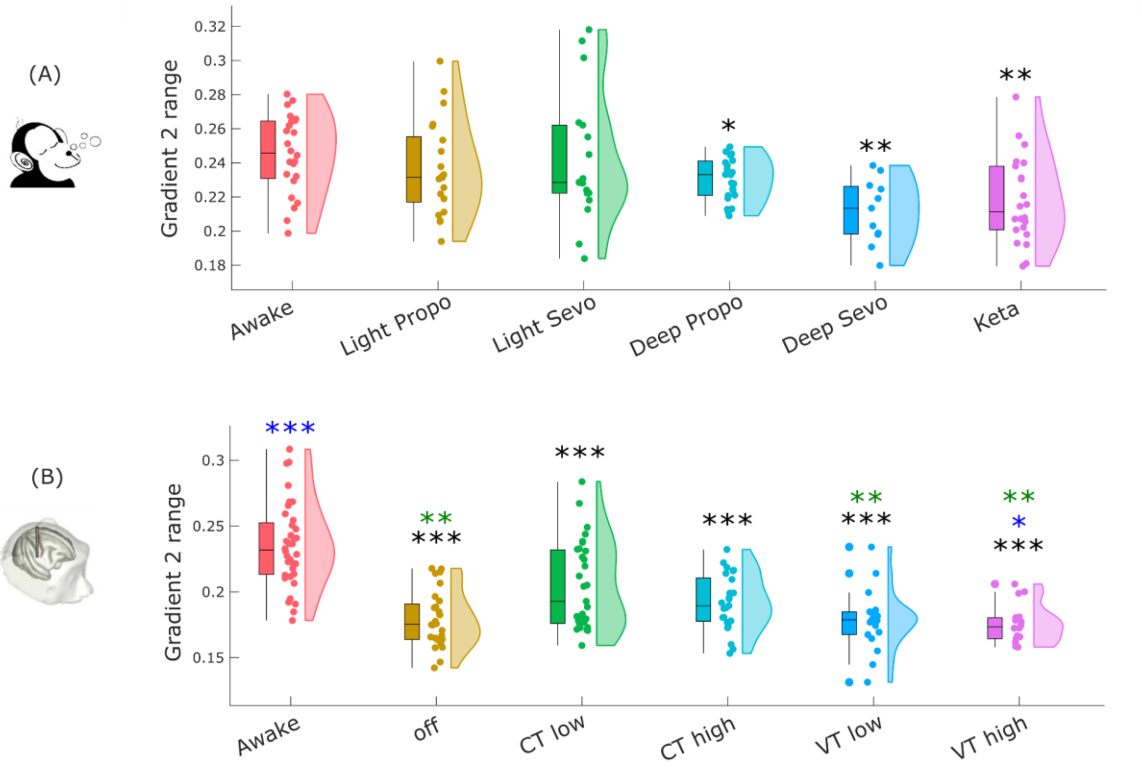


**Figure S2. Changes in the range of the second gradient of functional connectivity, across perturbations of consciousness.**. (A) Effect of different anaesthetics on the range of the second gradient of functional connectivity in macaques. (B) Effect of thalamic deep-brain stimulation on the range of the second gradient of functional connectivity in macaques during anaesthesia. Box plots indicate the median and interquartile range of the distribution. * $p < 0.05$; ** $p < 0.01$; *** $p < 0.001$ compared against Awake condition; * $p < 0.05$; ** $p < 0.01$; *** $p < 0.001$, compared against CT high condition; * $p < 0.05$; ** $p < 0.01$; *** $p < 0.001$, compared against CT low condition (FDR-corrected).

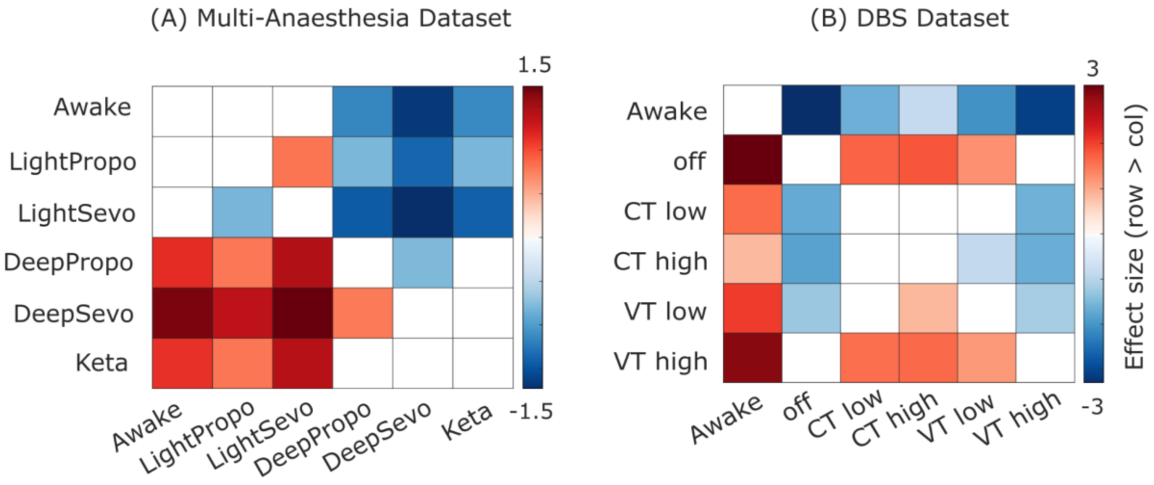


**Figure S3. Effect sizes for significant changes in harmonic energy across states of consciousness**. (A) Matrix shows the effect sizes for each statistical contrast between conditions of the multi-anaesthesia dataset; positive (negative) values indicate that the condition in the row is greater (lower) than the condition in the column. White entries indicate that the difference is not statistically significant. (B) Matrix shows the effect sizes for each statistical contrast between conditions in the DBS dataset; positive (negative) values indicate that the condition in the row is greater (lower) than the condition in the column. White entries indicate that the difference is not statistically significant.

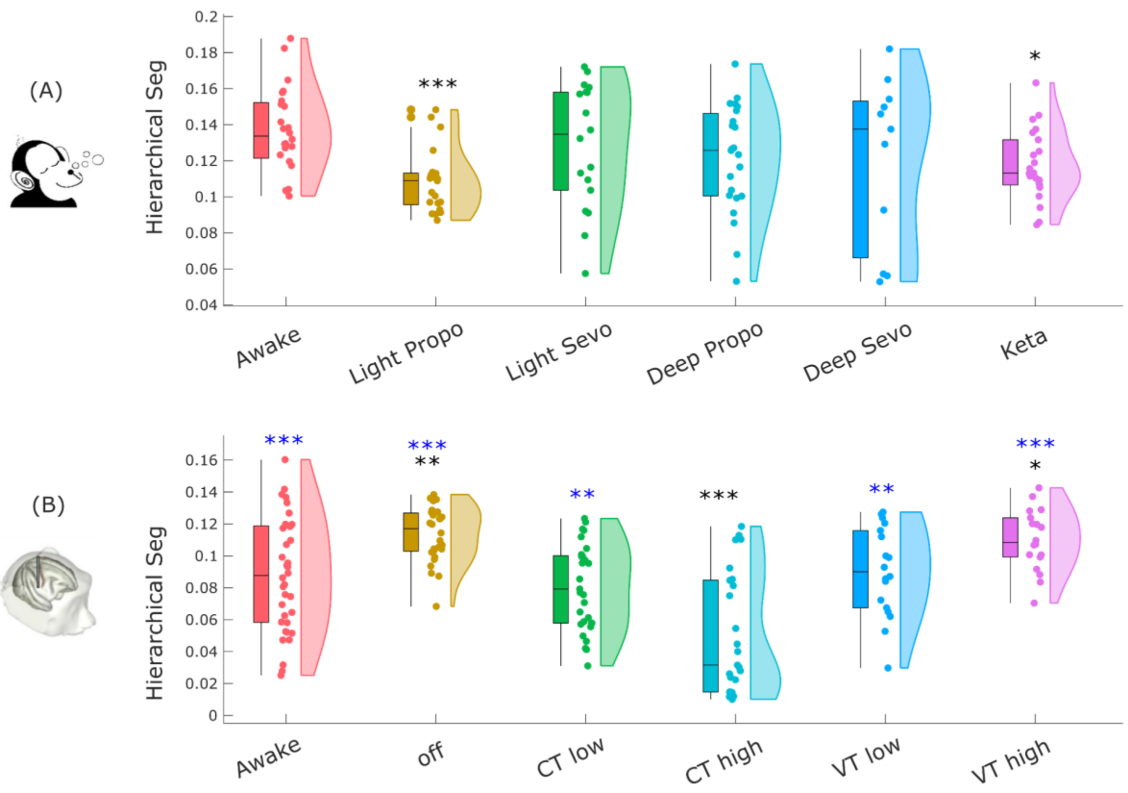


**Figure S4. Changes in eigenmode-based hierarchical segregation, across perturbations of consciousness.**. (A) Effect of different anaesthetics on the hierarchical segregation of functional eigenmodes. (B) Effect of thalamic deep-brain stimulation on the hierarchical segregation of functional eigenmodes during anaesthesia. Box plots indicate the median and interquartile range of the distribution. * $p < 0.05$; ** $p < 0.01$; *** $p < 0.001$ compared against Awake condition; * $p < 0.05$; ** $p < 0.01$; *** $p < 0.001$, compared against CT high condition; * $p < 0.05$; ** $p < 0.01$; *** $p < 0.001$, compared against CT low condition (FDR-corrected).

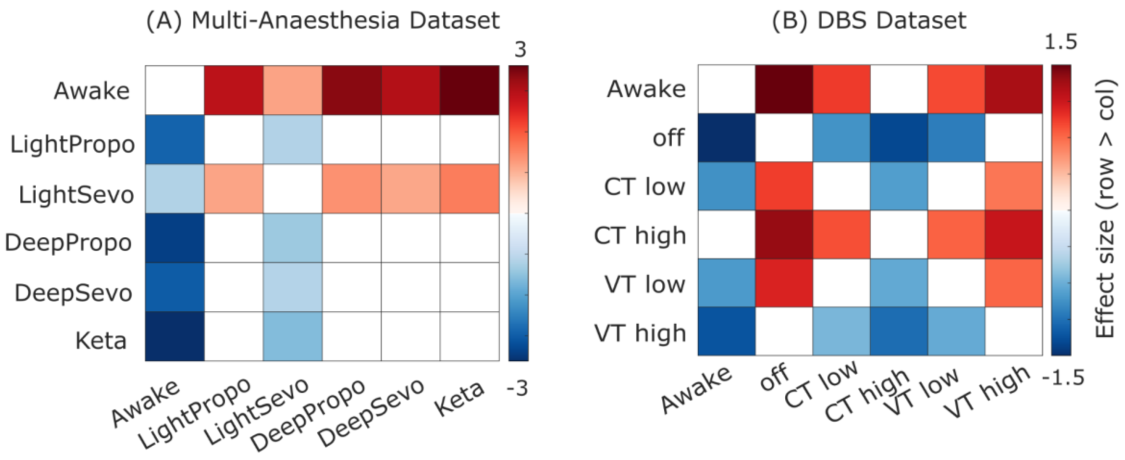


**Figure S5. Effect sizes for significant changes in hierarchical integration across states of consciousness**. (A) Matrix shows the effect sizes for each statistical contrast between conditions of the multi-anaesthesia dataset; positive (negative) values indicate that the condition in the row is greater (lower) than the condition in the column. White entries indicate that the difference is not statistically significant. (B) Matrix shows the effect sizes for each statistical contrast between conditions in the DBS dataset; positive (negative) values indicate that the condition in the row is greater (lower) than the condition in the column. White entries indicate that the difference is not statistically significant.
